# Novel and Reported Compensatory Mutations in *rpoABC* Associate Specifically with Predominant *Mycobacterium tuberculosis* Rifampicin Resistance Marker *rpoB*:S450L

**DOI:** 10.1101/2022.02.22.481565

**Authors:** D Conkle-Gutierrez, SM Ramirez-Busby, BM Gorman, A Elghraoui, S Hoffner, W Elmaraachli, F Valafar

## Abstract

**Background:** Rifampicin (RIF) is a key first-line drug used to treat tuberculosis, a pulmonary disease caused by *Mycobacterium tuberculosis*. However antibiotic resistance to RIF is prevalent despite an apparent fitness cost. RIF resistance is primarily caused by mutations in the RIF resistance determining region in the *rpoB* gene, at the cost of slower growth in rich media. Compensatory mutations in the genes *rpoA* and *rpoC* have been shown to alleviate this fitness cost. These compensatory mutations may explain how RIF resistant strains have spread so rapidly. However, the effect of compensation on transmission is still unclear, partly because of uncertainty over which *rpoABC* mutations compensate for which RIF resistance markers.

**Objectives:** We performed an association study on a globally representative set of 4309 whole genome sequenced clinical *M. tuberculosis* isolates to identify novel putative compensatory mutations, determine the prevalence of known and previously reported putative compensatory mutations, and determine which RIF resistance markers associate with these compensatory mutations.

**Results and Conclusions:** Only 20.0% (216/1079) of RIF resistant isolates carried previously reported high-probability compensatory mutations, suggesting existence of other compensatory mutations. Using a strict phylogenetic approach, we identified 18 novel putative compensatory mutations in *rpoC, rpoB*, and *rpoA*. Novel and previously reported compensatory mutations were strongly associated with the RIF^R^ marker *rpoB*:S450L, suggesting compensation may be specific to particular RIF^R^ markers. These findings will aid identification of RIF-resistant *M. tuberculosis* strains with restored fitness. Such strains pose a greater risk of causing resistant outbreaks.

## Introduction

Rifampicin (RIF) is an important first-line drug used to treat tuberculosis (TB), a pulmonary disease caused by *Mycobacterium tuberculosis*^1^. TB is a global pandemic, with an estimated 10 million cases and causing 1.2 million deaths in 2019^1^. Unfortunately, RIF resistance (RIF^R^) is prevalent. Half a million TB cases were RIF^R^ in 2019, including 3.3% of new TB cases and 17.7% of previously treated cases^1^. RIF^R^ and multidrug resistant (MDR) TB require 9 to 20 months of treatment with second-line drugs^1^.

RIF^R^ in *M. tuberculosis* is principally due to point mutations in the RIF resistance-determining region (RRDR)^2–4^, an 81-bp region (codons 426-452 in *M. tuberculosis,* codons 507-533 in *E. coli*) of the gene *rpoB^5^*, though two *rpoB* mutations outside the RRDR have also been shown to cause RIF resistance^6^. The *rpoB* gene encodes the β-subunit of RNA polymerase^5^. Mutations in the RRDR typically confer resistance by a change in the three-dimensional protein structure of the β-subunit, disrupting the RIF binding site^7^. The strong association between mutations in the RRDR and RIF resistance has led to the development of molecular tests and WHO recommended diagnostic platforms, including GeneXpert MTB/RIF, Truenat^8^, the line probe assays GenoType MTBDRplus VER 1 and 2 (Hain Lifescience, Germany), and Genoscholar NTM+MDRTB detection kit 2 (Nipro, Japan), and others^9–11^.

However, mutations in the RRDR have a fitness cost, slowing *M. tuberculosis* growth in vitro^12,13^ and in macrophage^12^. Despite the fitness cost, RIF^R^ strains continue to emerge, spread, and cause outbreaks^1^. This discrepancy may be due to compensatory mutations. Gagneux et al. observed that different *M. tuberculosis* strains with the same *rpoB* mutations had different fitness costs, and suggested the fitness costs were reduced in some strains by compensating mutations^13^. Later, Gagneux and Comas et al. identified 12 high-probability compensatory mutations (HCMs) in *rpoA* and *rpoC*, which encode the *β*’ and *α* subunits of RNA polymerase^14^. These 12 HCMs were carried by polyphyletic and exclusively RIF^R^ RRDR-variant *M. tuberculosis* isolates, suggesting these HCMs were selected for in RIF^R^ isolates^14^. Meanwhile in *Salmonella enterica*, several mutations in *rpoA* and *rpoC* have also been confirmed by mutagenesis to restore wild type growth in vitro to *rpoB* mutants^15^.

While compensatory mutations have been shown to increase the growth rate of RIF^R^ strains^14,15^ and prevent reversion to wildtype^15^, it is still unknown whether compensatory mutations increase the transmissibility of RIF^R^ tuberculosis. Several studies have shown the prevalence of putative compensatory mutations in clinical samples around the world^14,16–18^. However, a recent study found no association between putative compensatory mutations and transmission cluster size among MDR-TB strains^19^. A key challenge to answering this question is uncertainty over which *rpoABC* mutations compensate, and which RIF^R^ markers they compensate for.

In this work, we sought to expand the set of putative compensatory mutations and determine which specific RIF^R^ markers they associate with. We investigated the sequences of *rpoABC* genes in 4309 clinical *M. tuberculosis* isolates, of which 1079 were RIF^R^. This work identified 18 novel putative compensatory mutations. Additionally, both novel and previously reported putative compensatory mutations associated with the specific RIF^R^ maker *rpoB*:S450L, suggesting compensatory effects may be specific to particular *rpoB* mutations. These findings can help determine which RIF^R^ strains are compensated, which will both aid future studies in determining the role of compensation in transmission and eventually may serve as markers warning which resistant strains are most prone to causing outbreaks.

## Results

### Concordance of RIF^R^ genetic markers with phenotypic DST

We analyzed 4309 whole genome sequences and their phenotypic drug susceptibility testing (DST) results. In total 993 RIF^R^ and 78 RIF^S^ isolates (Tables 1, S1) harbored either a nonsynonymous mutation in the RRDR or one of the confirmed RIF^R^ conferring mutations outside the RRDR (*rpoB*:V170F or *rpoB*:I491F)^6^. Genotype-predicted RIF DST resulted in 92.0% sensitivity and 97.6% specificity. This was slightly lower than the sensitivity (93.8%, confidence interval 93.3%-94.2%) and specificity (98.2%, confidence interval 98.0%-98.3%) of prediction in a previous study of 27,063 isolates^6^. The lower specificity was likely due to the higher critical concentration use prior to 2021^20^. The high critical concentration has been shown to inconsistently classify RIF^R^ for isolates carrying any of six “borderline” mutations in the RRDR that confer lower level RIF^R^ (Table S2)^6,20,21^.

**Table 1.**
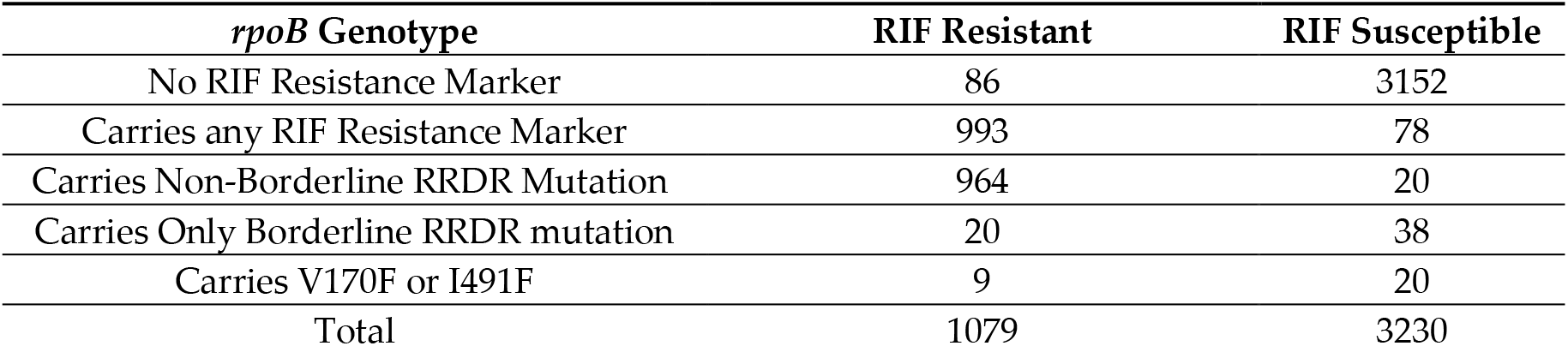
Number of rifampicin (RIF) resistant and RIF susceptible isolates carrying or not carrying RIF resistance markers. RIF resistance markers include non-borderline non-synonymous mutations in the rifampicin resistance-determining region (RRDR), borderline mutations in the RRDR, and the *rpoB* mutations V170F or I491F outside the RRDR.

The lower sensitivity may be from sampling bias favoring selection of discordant isolates for whole genome sequencing. There were 86 such discordant isolates (Table 1), with RIF^R^ DST results despite lacking any non-synonymous RRDR mutation (and lacking *rpoB*:V170F or *rpoB*:I491F). These discordant isolates could be the result of DST error, a resistant subpopulation, or an alternative mechanism of resistance. To find potential alternative mechanisms of resistance, the 86 discordant RIF^R^ isolates were queried for variants in *rpoB, rpoC*, and *rpoA*. Among 42 of the 86 discordant RIF^R^ isolates were 17 unique mutations, of which 7 were exclusive to RIF^R^ isolates (Table S3). In the remaining 44 discordant RIF^R^ isolates there were no mutations in *rpoA*, *rpoB*, or *rpoC*.

### High-probability compensatory mutations associated with the common RIF^R^ marker rpoB:S450L

Comas et al. previously identified 12 HCMs in *rpoA* and *rpoC* that likely compensated for the fitness cost of RRDR mutations in vitro^14^. We searched for these HCMs in the 4309 isolates (Table 2). Of the 217 isolates carrying HCMs, all but one were RIF^R^. The RIF^S^ isolate carried the common RIF^R^ conferring RRDR mutation *rpoB*:S450L and the HCM *rpoC*:V483G (Table S1). The RIF^S^ DST result in this isolate was likely laboratory error, as the isolate carried known markers for multiple drugs despite pan susceptible DST results.

**Table 2.**
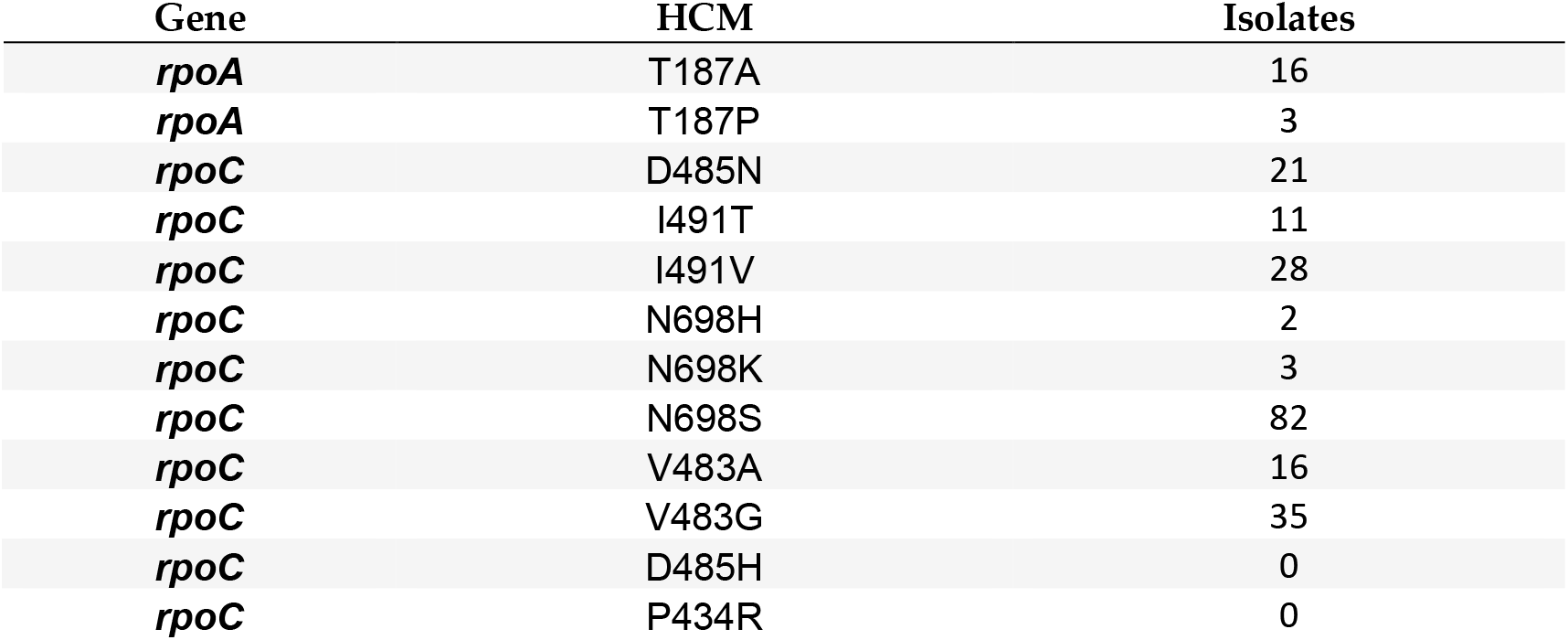
Number of isolates carrying each of the 12 previously identified^14^ high-probability compensatory mutations (HCMs) in *rpoA* and *rpoC*.

Isolates carrying *rpoB*:S450L were 48.8 times more likely to carry HCMs than isolates carrying other RIF^R^ markers (Table 3, odds ratio = 48.8, Fisher’s exact test p = 7.37e-28). Only two isolates carried an HCM and lacked *rpoB*:S450L. One such isolate carried the non-borderline RRDR mutation *rpoB*:Q432P and the HCM *rpoC*:V483G (Table S1). The other isolate carried the RIF^R^ marker *rpoB*:V170F and carried the HCM *rpoA*:T187P (Table S1). This isolate also potentially included a subpopulation carrying *rpoB*:S450L. The *rpoB*:S450L variant was supported in the isolate by 7 of the 78 reads mapped to this locus. However, it is uncertain whether these sequencing reads was the result of a genuine subpopulation or sequencing error.

**Table 3.** Number of isolates carrying previously identified^14^ high-probability compensatory mutations (HCMs) and the common RIF^R^ conferring mutation *rpoB*:S450L. Only isolates carrying RIF^R^ markers are included in these counts.

| Compensation Status | Carries *rpoB*:S450L | No *rpoB*:S450L |
| --- | --- | --- |
| Carries HCM | 215 | 2 |
| No HCM | 587 | 267 |

### Previously reported putative compensatory mutations

Only 20.0% (216/1079) of RIF^R^ isolates carried an HCM. We then queried for 97 previously reported putative compensatory mutations from their study and others^14,16,17,19,22–24^. Out of these 97 previously reported mutations, 73 were carried by at least one of the 4309 isolates. In total 523 RIF^R^ and 4 RIF^S^ isolates each carried at least one of these previously reported mutations (Table S4). Only 56.3% (608/1079) of RIF^R^ isolates carried HCMs or previously reported putative compensatory mutations.

### Novel putative compensatory mutations

We then searched for novel putative compensatory mutations in *rpoABC* with the following criteria: i) the mutation must be carried by at least one RIF^R^ isolate lacking an HCM^14^; ii) the mutation must not be carried by any RIF^S^ isolate; iii) the mutation must be carried by at least two isolates. We then determined whether each novel putative compensatory mutation was carried by a polyphyletic group of isolates, using a phylogenetic tree (Figure 1). These criteria were developed based on the initial criteria set by Comas et al.^14^ to discover candidate variants, later built upon in Liu et al.^19^, and here modified to use a phylogenetic tree to define relationships among the isolates rather than SNP distance.

**Figure 1.**
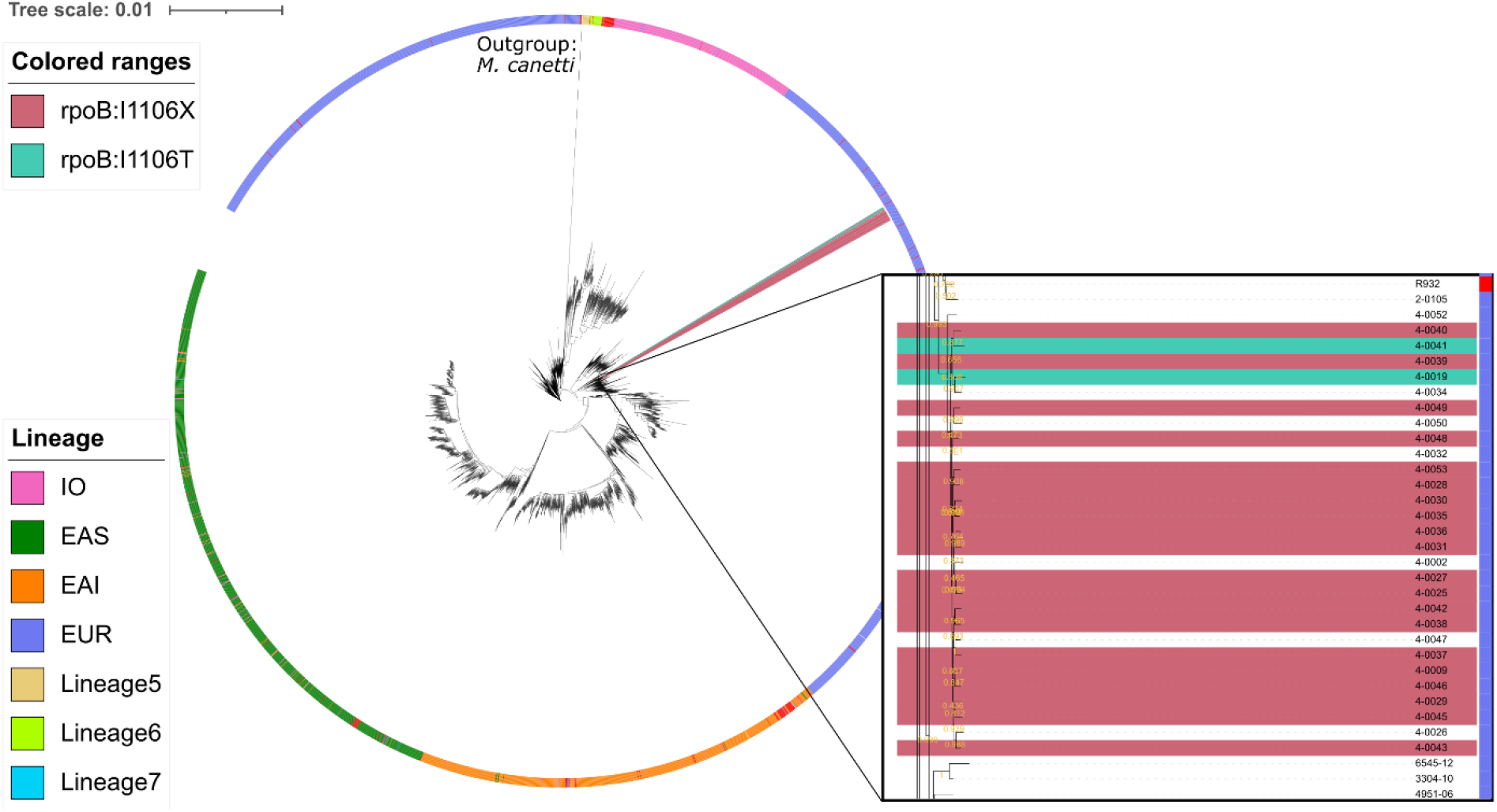
Maximum likelihood phylogenetic tree from whole-genome SNPs using FastTree of the 4309 *M. tuberculosis* complex isolates with *M. canettii* (NC_019951.1) as the outgroup. In the outer ring, colors represent lineages that were determined by MIRU-VNTR/spoligotyping. The colored wedge highlights the 22 isolates with the SNP *rpoB*:I1106T and frameshift *rpoB*:I1106X. The smallest monophyletic group containing all 22 isolates includes an additional 7 isolates. All 29 isolates in the group were collected from the National Health Laboratory Service of South Africa in Johannesburg, South Africa.

These filters identified 18 novel putative compensatory mutations in 68 isolates, of which 14 mutations were polyphyletic (Table 4). Together 62.1% (670/1079) of RIF^R^ isolates carried either an HCM, a previously reported putative compensatory mutation, or a novel putative compensatory mutation. This method also independently corroborated 29 previously reported putative compensatory mutations (Table S5), including *rpoB*:I480V which was recently shown to compensate in *M. smegmatis*^24^. That mutations identified as compensatory by this method were independently reported in contemporary studies supports the reproducibility and validity of the method.

**Table 4.** Number of isolates carrying novel putative compensatory mutations in *rpoA* and *rpoC*. The mutations carried by a monophyletic group of isolates are indicated by an asterisk (*) next to their number of isolates. All other mutations were each carried by polyphyletic groups of isolates. Each reported mutation was carried by at least two isolates, was carried exclusively by RIF^R^ isolates, and was carried by at least one RIF^R^ isolate lacking a previously identified^14^ high-probability compensatory mutation.

| Gene | Novel Putative<br>Compensatory Mutation | Isolates |
| --- | --- | --- |
| *rpoB* | I1106X | 20 |
| *rpoC* | R770H | 8 |
| *rpoA* | E184D | 6 |
| *rpoA* | A180V | 3 |
| *rpoA* | GC-63G | 3 |
| *rpoB* | R827H | 3 |
| *rpoB* | Y564H | 3 |
| *rpoC* | D943G | 3 |
| *rpoC* | L402F | 3* |
| *rpoC* | F831L | 3 |
| *rpoA* | CGGG-84C | 2 |
| *rpoB* | I1106T | 2 |
| *rpoB* | K1102T | 2* |
| *rpoC* | E1033A | 2 |
| *rpoC* | E750A | 2 |
| *rpoC* | H748P | 2* |
| *rpoC* | S428A | 2 |
| *rpoC* | S838C | 2 |

As with the previously reported mutations, most novel putative compensatory mutations carried *rpoB*:S450L (Table 5). Isolates carrying *rpoB*:S450L were 18.4 times more likely to carry an HCM or putative compensatory mutation than isolates carrying other RIF^R^ markers (Table S6, odds ratio = 18.4, Fisher’s exact test p = 1.18e-74).

**Table 5.** Number of isolates carrying rifampicin resistance markers common in the dataset. All RRDR mutations carried by at least 20 isolates are included, as well as the known rifampicin resistance marker *rpoB*:I491F. Columns “*R*” and “S” report the number of rifampicin resistant and susceptible isolates carrying each mutation, according to phenotypic drug susceptibility testing. Column “HCM” reports the number of isolates carrying any of the 12 high-probability compensatory mutations previously reported by Comes et al.^14^. Column “Reported Putative Comp” reports the number of isolates carrying both the indicated resistance marker and any previously reported^14,16,17,19,22,23^ putative compensatory mutations. Column “Novel Putative Comp” reports the number of isolates carrying both the indicated resistance marker and any of the novel putative compensatory mutations identified in this study. The last five rows report the number of isolates carrying combinations of multiple of the markers from the table. Combinations carried by zero isolates are excluded.

| Marker | R | S | HCM | Reported Putative Comp | Novel Putative Comp |
| --- | --- | --- | --- | --- | --- |
| S450L | 801 | 1 | 215 | 507 | 42 |
| D435V | 40 | 2 | 1 | 2 | 1 |
| L452P | 32 | 10 | 0 | 2 | 17 |
| H445Y | 31 | 2 | 0 | 1 | 0 |
| D435G | 24 | 1 | 0 | 0 | 17 |
| H445D | 21 | 1 | 0 | 1 | 0 |
| I491F | 6 | 20 | 0 | 1 | 0 |
| S450L D435V | 3 | 0 | 1 | 2 | 0 |
| S450L H445Y | 1 | 0 | 0 | 1 | 0 |
| S450L H445D | 1 | 0 | 0 | 0 | 0 |
| L452P H445Y | 0 | 1 | 0 | 0 | 0 |
| L452P D435G | 16 | 0 | 0 | 0 | 13 |

This method classified five frameshift causing single base deletions as novel putative compensatory mutations (*rpoB*:D435X, *rpoB*:I1106X, *rpoC*:E1137X, *rpoC*:P390X, *rpoC*:V483X), four of which we excluded as likely sequencing error. However, *rpoB*:I1106X, carried by 20 closely related isolates (Figure 1), was unlikely to be a random error. The SNP *rpoB*:I1106T was also carried by two related isolates (Figure 1), suggesting that *rpoB*:I1106X may be a systematically missed variant call of the same underlying mutation. At the nucleotide level, the SNP *rpoB*:I1106T (T3317C) extended a homopolymer (from ATCCCG to ACCCCG). The 20 isolates with *rpoB*:I1106X were SMRT sequenced with P4C2 chemistry and variant called with PBHoover^25^, while the two isolates with *rpoB*:I1106T were SMRT sequenced with P6C4 chemistry and variant called with mummer^26^ after de novo assembly.

The 22 isolates with *rpoB*:I1106T/X were all RIF^R^ and each carried two RRDR mutations at the codons 452 and 435 (*rpoB*:D435G/X and *rpoB*:L452P/X). These rare RRDR mutations were carried in only 33 isolates. Isolates carrying *rpoB*:I1106T/X were at least 663 times more likely to carry either *rpoB*:D435G/X or *rpoB*:L452P/X than isolates without *rpoB*:I1106T/X (Table S7, odds ratio 95% confidence interval: 663 to ∞, p-value = 1.71e-44).

HCMs and putative compensatory mutations in *rpoA* were exclusively in the RNA polymerase Rpb3/Rpb11 dimerization domain (Figure 2A). In *rpoC* the HCMs and putative compensatory mutations primarily clustered in Domain 2 (Figure 2C).

**Figure 2.**
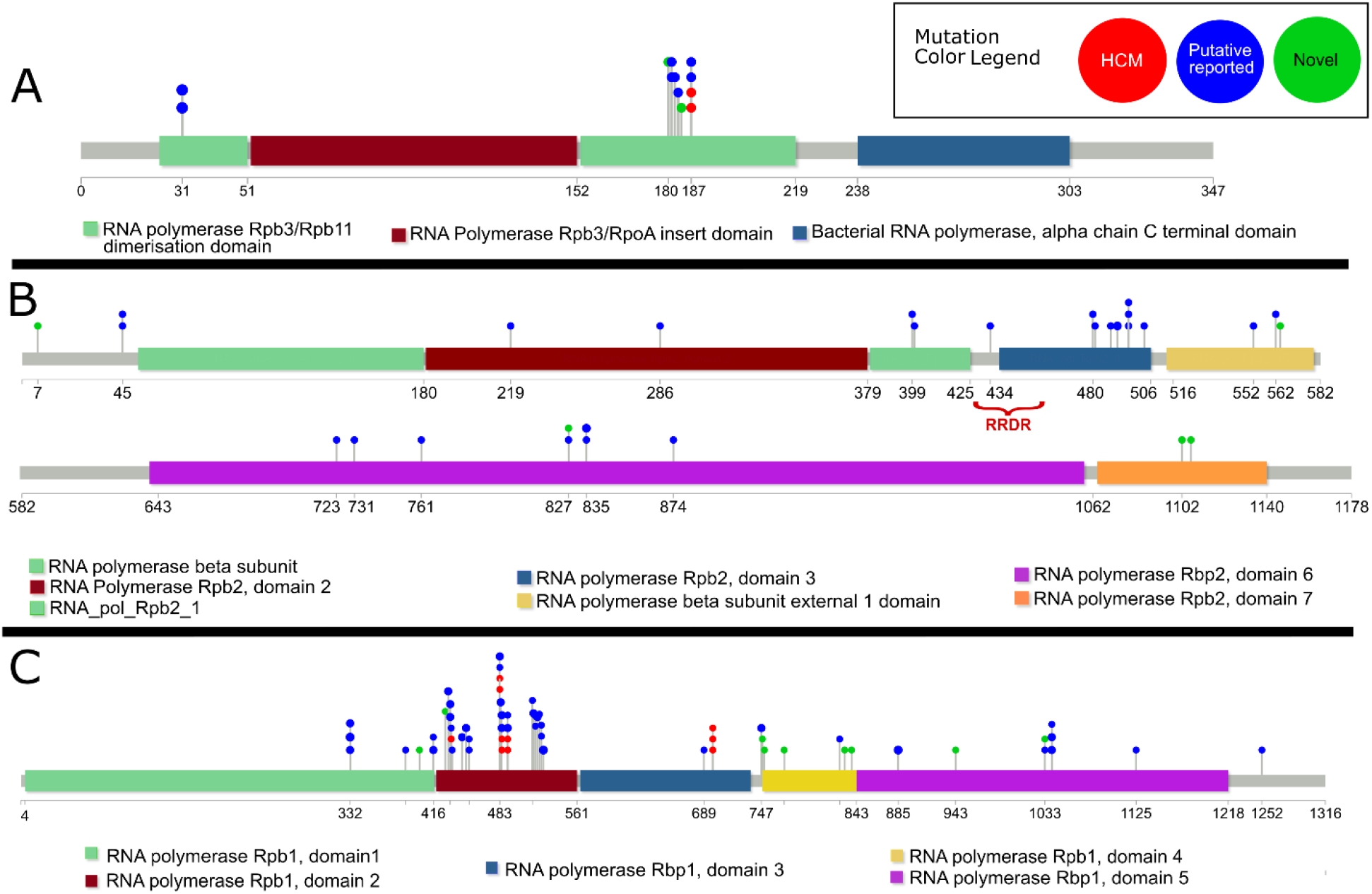
Lollipop diagram of previously reported^14^ high-probability compensatory mutations (“HCM”), previously reported^14,16,17,19,22–24^ putative compensatory mutations (“Putative reported”), and novel putative compensatory mutations (“Novel”) in the *M. tuberculosis* genes A) *rpoA*, B) *rpoB*, and C) *rpoC*. The functional domains of each gene are indicated, as well as the rifampicin resistance determining region (RRDR) in *rpoB*. Only previously reported putative compensatory mutations carried by isolates in this dataset are included.

## Discussion

Resistance to RIF is prevalent despite the fitness cost of RIF^R^ conferring mutations. Multiple RIF^R^ mutations in *rpoB* have been shown to reduce growth rate in vitro^13,14^ and in macrophage^12^. Yet rifampicin resistant strains continue to spread, and in some countries constitutes an increasing threat to an effective TB control^1^. One potential explanation for this discrepancy is compensatory mutations in *rpoA, rpoB,* and *rpoC*. Three mutations in these genes have been observed in clinical RIF^R^ *M. tuberculosis* isolates with a restored wildtype growth rate in vitro^14^. Many more mutations in these genes have associated with clinical RIF^R^ strains^14,16,17,19^. And five mutations in *rpoB* have recently been confirmed by mutagenesis and competitive fitness assays to compensate for *rpoB*:S450L in *M. smegmatis*^24^. RIF^R^ strains with compensatory mutations may have higher transmission rates than RIF^R^ strains without them^16,18,23,27^, though this is disputed^19^. Determining the effect of compensatory mutations on transmission is difficult because the full set of compensatory mutations is still unknown, making it uncertain which transmission clusters have compensatory mutations. We expanded the set of putative compensatory mutations by examining *rpoABC* mutations in 4309 whole genome sequenced clinical *M. tuberculosis* isolates. We also uncovered evidence that each compensatory mutation may only compensate for a specific RIF^R^ mutation, rather than all compensatory mutations compensating for all RIF^R^ mutations. If confirmed, these finding will improve accuracy in determining which transmission clusters are compensated.

Previously reported compensatory mutations^14,16,17,19,22–24^ accounted for only 56.3% (608/1079) of RIF^R^ isolates. The remaining RIF^R^ isolates either lacked compensation or carried novel compensatory mutations. Using strict criteria, we identified 18 novel putative compensatory mutations in 68 of these isolates (Table 4). Our criteria additionally independently identified 29 of the previously reported putative compensatory mutations (Table S5). One of these independently identified mutations, *rpoB*:I480V, was confirmed by mutagenesis to compensate for *rpoB*:S450L in vitro in M. smegmatis^24^. This independent corroboration of 29 putative compensatory mutations identified by our method supports the validity of the 18 novel putative compensatory mutations identified by our criteria. These 18 novel putative compensatory mutations, together with those previously reported, may aid future molecular epidemiological studies identify transmission clusters carrying compensatory mutations. If compensatory mutations do increase transmission rates of RIF^R^ strains, then these mutations may eventually serve as markers to warn clinicians which RIF^R^ strains are more likely to cause outbreaks.

Another potential barrier to identifying compensated strains is the specificity of compensatory mutations. Only three RIF^R^ markers associated with compensatory mutations: *rpoB*:S450L, *rpoB*:L452P, and *rpoB*:D435G (Tables 3, 5, S5, and S7). The novel putative compensatory mutation *rpoB*:I1106T was specific to isolates with a pair of rare RRDR mutations, *rpoB*:L452P and *rpoB*:D435G. Most other HCMs and putative compensatory mutations associated with the prevalent RRDR mutation *rpoB*:S450L. This is consistent with recent studies showing associations between *rpoB*:S450L and compensatory *rpoC*^28^ and *rpoB*^24^ mutations. It is possible that compensatory mutations each only compensate for specific RRDR mutations. If so, predicting which RIF^R^ strains will likely cause outbreaks requires not only a catalogue of compensatory mutations, but knowledge of which combinations of *rpoABC* mutations provide a compensatory effect.

This mutation-specific-compensation would potentially help explain the high prevalence of *rpoB*:S450L among RIF^R^ isolates. A recent study of 27,063 isolates found *rpoB*:S450L in 66.2% of RIF^R^ isolates (6536/9869)^6^. If most compensatory mutations only compensate for *rpoB*:S450L, then RIF^R^ strains with this mutation would be more likely to develop compensation and increase their fitness. However, the high prevalence of *rpoB*:S450L compared to other RIF^R^ markers could also be due to its overall lower fitness cost, rather than specific compensation. In previous competitive fitness assays, *rpoB*:S450L had a lower fitness cost then other RIF^R^ mutations, regardless of genetic background (though *rpoB*:S450L still caused slower growth than wild type unless paired with a compensatory mutation)^13^. Furthermore *rpoB*:S450L is the most common mutation among in vitro selected RIF^R^ mutants^29^. Indeed, Pengjiao Ma et al. have suggested that this lower base fitness cost is not only responsible for the higher prevalence of *rpoB*:S450L, but also its association with compensatory mutations^24^. Pengjiao Ma et al. have suggested that the lower base fitness cost of *rpoB*:S450L gives isolates with it more time to gain compensatory mutations^24^.

To determine whether the association between *rpoB*:S450L and compensatory mutations is due to mutation-specific-compensation or due to other factors such as the lower base fitness cost of *rpoB*:S450L, further competitive fitness studies are needed. Currently *rpoB*:S450L is the only RIF^R^ marker with in vitro evidence in *M. tuberculosis*^14^ or *M. smegmatis*^24^ that its fitness cost is reduced by compensatory mutations, though another RIF^R^ conferring mutation has shown compensation in *Salmonella enterica*^15^. Further competitive fitness studies are needed to test whether compensatory mutations restore fitness to other RIF^R^ markers. Testing these hypotheses is important for identifying compensated RIF^R^ strains, which may be at greater risk of causing outbreaks.

Beyond *rpoB*:S450L, compensatory mutations may enable rarer RIF^R^ markers to overcome their greater fitness cost and spread, causing RIF^R^ outbreaks. This may have been the case with the 22 closely related RIF^R^ isolates carrying the novel putative compensatory mutations *rpoB*:I1106T/X and the rare RRDR mutations L452P/X and D435G/X, collected in Johannesburg, South Africa. However, as *rpoB*:I1106T/X were only carried by isolates from this cluster, it is uncertain whether the mutation truly improved fitness, or was simply a neutral mutation carried by the common ancestor of the outbreak. Monophyletic mutations can still effect phenotype^30^, however as with any association, follow-up mutagenesis is needed to show causation.

This study had several limitations. The study combined sets of isolates sequenced by different platforms, each with different sequencing biases. For example, the RS1 SMRT sequenced isolates had lower than average coverage, and the Illumina sequenced isolates have known difficulties with homopolymers and high GC regions^31^. Additionally, the novel putative compensatory mutations identified in this study were implicated only by association and require future mutagenesis experiments to confirm whether they truly compensate for fitness costs. Finally, the criteria to identify these novel putative compensatory mutations was conservative and thus our expanded set of compensatory mutations is more reliable, but not comprehensive. Further study is needed to determine the full set of compensatory mutations.

This study identified 18 novel *rpoABC* mutations that putatively compensate for the fitness cost of *rpoB*:S450L. These mutations may aid future investigation of the effect of compensatory mutations on RIF^R^ TB strain transmission, and eventually aid the detection of strains at high risk of causing RIF^R^ outbreaks. This study additionally found that compensatory mutations associated with specific RIF^R^ markers, highlighting the need for future study of which RIF^R^ markers are prone to compensation.

## Materials and Methods

### Sample Selection

We analyzed a total of 4309 whole genome sequences. Of these 314 were sequenced on Pacific Bioscience’s (PacBio) Real-time Sequencer (RS) and RS II platforms (Bioproject: PRJNA353873). These isolates originated from Hinduja National Hospital (PDHNH) in Mumbai, India; the Phthisiopneumology Institute (PPI) in Chisinau, Moldova; the Tropical Disease Foundation (TDF) in Manila, Philippines; the National Health Laboratory Service of South Africa (NHLS) in Johannesburg, South Africa.

We also downloaded 3995 whole genome sequences from NCBI’s Sequence Read Archive (SRA) database using SRA Toolkit’s fastqdump^32^ (Bioprojects: PRJEB2221, PRJEB5162, PRJEB6276, PRJEB7281, PRJEB7727, PRJEB9680, PRJNA282721, PRJEB2138). All downloaded raw reads were sequenced on Illumina platforms. These genomes were isolated from patients originating from the UK, Sierra Leon, South Africa, Germany, Uzbekistan^33^, and Russia^22,23,34^.

### Drug Susceptibility Testing

The RIF susceptibility testing for the PacBio and Illumina sequenced isolates were described in^22,23,33–36^, respectively. Briefly, all samples were tested on the BACTEC MGIT960 platform and a rifampicin (RIF) critical concentration of 1μg/ml.

### Whole Genome Sequencing

The DNA sequencing protocol for the PacBio RS and RSII platforms was described previously^37^. The sequencing protocol for genomes sequenced on Illumina Genome Analyzer, MiSeq, or HiSeq platforms were described previously^22,23,33,34^.

### Alignment and Variant Calling

For PacBio sequences, raw reads were aligned to *M. tuberculosis* H37Rv reference strain (Genbank accession number NC_000962.3) utilizing SMRT Analysis’ Basic Local Alignment with Successive Refinement (BLASR) with default parameters (v1.3)^38^. For variant calling, a custom software— PBHoover^25^ (manuscript submitted)—corrected aligned reads and called variants based on a maximum likelihood criterion.

For 46 PacBio isolates that passed assembly QC, whole genomes were assembled with HGAP as described previously^39^, then aligned to H37Rv using dnadiff (v1.3) from the mummer suite^26^. Variants were converted to VCF format using a custom script, mummer-snps2vcf (https://gitlab.com/LPCDRP/mummer-extras/-/blob/master/src/mummer-snps2vcf).

All Illumina short-read data were processed as follows: trimmomatic (v0.36)^40^ trimmed adapters from raw reads; bowtie2 (v2.2.4)^41^ aligned reads to the *M. tuberculosis* H37Rv reference sequence (Genbank accession NC_000962.3); SAMtools (v1.3.1)^42^ sorted, filtered out reads with a mapping quality of less than 20, and created an mpileup file for each isolate; VarScan2 (v2.3)^43^ called and filtered variants with a minimum quality of 20, a minimum depth of 10, and strand filter set to false.

For two Illumina sequenced isolates carrying HCMs and not carrying variant *rpoB*:S450L, LoFreq^44^ was used to check for potential subpopulations with that variant by calculating the fraction reads supporting *rpoB*:S450L as a minor variant.

VCF formatted files were further annotated with Variant Effect Predictor (VEP) (v87)^45^ to determine the consequence of each variant.

### Lineage Determination

For isolates from Bioproject PRJNA353873, we utilized previously determined MIRU-VNTR and spoligotyping patterns^36^ and determined lineage using the TB-Insight webserver^46^.

For all remaining isolates, we de novo assembled each genome using SPAdes^47^ (v3.9.0) with default parameters. A custom Python script, MiruHero (https://gitlab.com/LPCDRP/miru-hero), queried each draft genome for 12 MIRUs and all spacers to determine the MIRU-VNTR and spoligotyping pattern. MiruHero implements the lineage by rules criteria developed for TB-Insight^48^ to determine lineage.

### Phylogeny

We constructed a phylogenetic tree that included all 4309 isolates. A custom Python script parsed all VCF files to determine all positions that contained SNPs across the population. For each position, the base call was retrieved (either variant or reference) and recorded for each isolate. *Mycobacterium canettii* (NC_01995.1) was chosen as the outgroup. To identify base calls for the SNP positions determined previously in each outgroup, we used progressiveMauve^49^ and wrote a custom Python script that identified the base calls in the outgroups for each SNP position, missing positions were populated with N. The resulting multisequence FASTA contained sequences with the same length, therefore, no alignment was necessary. FastTree (v2.1.10) was used to generate a maximum likelihood tree using default parameters. The tree was visualized with the Interactive Tree of Life^50^ (iToL), coloring was determined by lineage (see *Lineage Determination* above).

### Identifying Putative Compensatory Mutations

We identified putative compensatory mutations in *rpoA*, *rpoB*, and *rpoC* using the following criteria: i) the mutation must be carried by at least one RIF^R^ isolate lacking an HCM^14^; ii) the mutation must not be carried by any RIF^S^ isolate; iii) the mutation must be carried by at least two isolates (as mutations carried by multiple isolates are more likely to have a beneficial effect on fitness). Three frameshift causing single base deletions were also excluded. We then determined whether each novel putative compensatory mutation was carried by a polythetic group of isolates using the phylogenetic tree and ETE Toolkit v3.1.1^51^.

HCMs, previously reported putative compensatory, and novel compensatory mutations were mapped to the *rpoA, rpoB*, and *rpoC* genes with known domains using Lollipop^52^, with manual adjustments in Inkscape to colors and the heights of mutation labels.

## Data Availability

All supplemental tables are available at https://doi.org/10.5281/zenodo.6082557.

## Acknowledgements

We would like to acknowledge James O’Neill from the Laboratory for Pathogenesis of Clinical Drug Resistance and Persistence (LPCDRP) his role in the analysis of compensatory mutations outside of *rpoC* and *rpoA* and background research on drug susceptibility accuracy and Scott Mitchell from the LPCDRP for assistance in creating the phylogenetic tree.

## Transparency Declarations

Authors have no conflict of interest to declare in this project. Other than the funding declared in the “Funding Information” section, no other funding has been received by the authors for this project.

